# Temporal discrimination from the interaction between dynamic synapses and intrinsic subthreshold oscillations

**DOI:** 10.1101/727735

**Authors:** Joaquin J. Torres, Fabiano Baroni, Roberto Latorre, Pablo Varona

## Abstract

The interaction between synaptic and intrinsic dynamics can efficiently shape neuronal input-output relationships in response to temporally structured spike trains. We use a neuron model with subthreshold oscillations receiving inputs through a synapse with short-term depression and facilitation to show that the combination of intrinsic subthreshold and synaptic dynamics leads to channel-specific nontrivial responses and recognition of specific temporal structures. We employ the Generalized Integrate-and-Fire (GIF) model, which can be subjected to analytical characterization. We map the temporal structure of spike input trains to the type of spike response, and show how the emergence of nontrivial input-output preferences is modulated by intrinsic and synaptic parameters in a synergistic manner. We demonstrate that these temporal input discrimination properties are robust to noise and to variations in synaptic strength, suggesting that they likely contribute to neuronal computation in biological circuits. Furthermore, we also illustrate the presence of these input-output relationships in conductance-based models.

**Author summary:** Neuronal subthreshold oscillations underlie key aspects of information processing in single neuron and network dynamics. Dynamic synapses provide a channel-specific temporal modulation of input information. We combine a neuron model that displays subthreshold oscillations and a dynamic synapse to analytically assess their interplay in processing trains of spike-mediated synaptic currents. Our results show that the co-action of intrinsic and synaptic dynamics builds nontrivial input-output relationships, which are resistant to noise and to changes in synaptic strength. The discrimination of a precise temporal structure of the input signal is shaped as a function of the joint interaction of intrinsic oscillations and synaptic dynamics. This interaction can result in channel-specific recognition of precise temporal patterns, hence greatly expanding the flexibility and complexity in information processing achievable by individual neurons with respect to temporal discrimination mechanisms based on intrinsic neuronal dynamics alone.

## Introduction

Neuronal subthreshold oscillations underlie key mechanisms of information discrimination in single cells [1–4] while dynamic synapses provide channel-specific input modulation [5–9]. Previous studies have shown that intrinsic neuronal properties, in particular subthreshold oscillations, constitute a biophysical mechanism for the emergence of nontrivial single-cell input-output preferences, e.g., preference towards decelerating vs. accelerating input trains of the same total duration and constituent interspike intervals (ISIs) [10, 11]. It has also been shown that short-term synaptic dynamics, in the form of short-term depression and/or short-term facilitation, can provide a channel-specific mechanism for the enhancement of the postsynaptic effects of temporally specific input sequences [8, 12], suggesting a functional role for the multiplexing of information in neuronal circuits [13]. This adds to other tuning properties of synaptic integration [14]. While intrinsic oscillations and synaptic dynamics are typically studied independently, it is reasonable to hypothesize that their interplay can lead to more selective and complex temporal input processing.

Here, we present an analytical study on the interaction between subthreshold oscillations and short-term synaptic dynamics. We investigated whether and under which conditions the combination of intrinsic subthreshold oscillations and short-term synaptic dynamics synergistically enables the emergence of robust and channel-specific selectivity of temporal structure in neuronal input-output transformations. We calculated analytically the voltage trajectories and spike output of Generalized Integrate-and-Fire (GIF) model neurons in response to temporally distinct trains of input spikes delivered through a dynamic synapse. In particular, we considered triplets of excitatory postsynaptic currents (EPSCs) in a range that covers intrinsic and synaptic time scales, and analyzed the model output as intrinsic and synaptic parameters were varied.

Our results show that intrinsic and synaptic dynamics interact in a complex manner for the emergence of specific input-output transformations resulting from the temporal discrimination of input trains. In particular, localized nontrivial preferences emerge from intrinsic and synaptic dynamics with coincident temporal preferences, while more complex and distributed selectivity can be observed for intrinsic and synaptic dynamics with mismatched temporal preferences. Throughout our analysis, we discuss the conditions for robustness of the observed input-output relationships, which we show that appear also in conductance-based models. The analysis presented in this paper generalizes and extends our previous computational study on the action of a depressing synapse on a neuronal model with subthreshold oscillations in response to bursting input [15]. The results described below show that combining synaptic facilitation and depression with intrinsic subthreshold oscillations can result in selective and nontrivial recognition of distinct input spike trains, hence constituting a channel-specific mechanism for the emergence of selective neuronal responses.

This work is motivated by experimental evidence describing distinct short-term synaptic dynamics in different afferents projecting to the same area and/or neuronal population. Variability in short-term synaptic dynamics have been described in several systems, including variability between thalamocortical connections to the visual and somatosensory cortices [16, 17], between visceral afferents from the brainstem to the dorsal vagal complex [18], and between hindbrain afferents to the midbrain torus semicircularis (analogous to the mammalian inferior colliculus) in the electrosensory system in weakly electric fish [19]. In the case of distinct vestibular and visual pathways to the cerebellum, variability in short-term synaptic dynamics have been demonstrated at the single postsynaptic neuron level [20]. However, the functional consequences of the variability in short-term synaptic dynamics, and of the interaction between synaptic dynamics and intrinsic subthreshold oscillations, are still unexplored. Here, we advance specific hypotheses that link heterogeneous synaptic dynamics and subthreshold oscillations to their distinct computational function in the framework of input-output temporal pattern transformation of spiking activity. More generally, we discuss the impact of single-channel/single-neuron temporal input discrimination in the context of information processing based on heterogeneous elements.

## Methods

### Neuron Model

The use of an analytically tractable model to assess the interaction between intrinsic subthreshold oscillations and dynamic synapses provides rigorous theoretical predictions regarding the synergistic temporal co-action of synaptic and intrinsic dynamics. We considered the Generalized Integrate-and-Fire (GIF) neuron as a minimal and analytically tractable model that exhibits subthreshold damped oscillations [21–23]. The GIF model is defined as

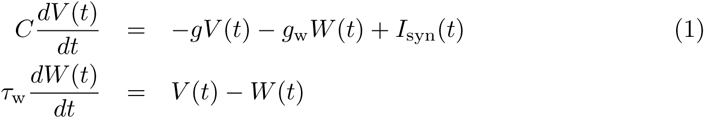

where *V* represents the membrane potential deviation from the leak reversal potential and *W* is a gating variable characterizing the membrane dynamics, which results from the linearization of voltage-gated ionic currents [21, 24]. *C* is the membrane capacitance, *g* is the effective leak conductance, and *g*_w_ and *τ*_w_ are the effective ionic conductance and time constant associated with the *W* variable, which corresponds to the activation of a restorative intrinsic current *I*_w_ = *g*_w_*W* (*t*). *I*_syn_ represents the synaptic current resulting from action potential generation in presynaptic neurons. The parameter values of the canonical GIF model that we considered for most of our results are reported in Table 1. The model can be rewritten as:

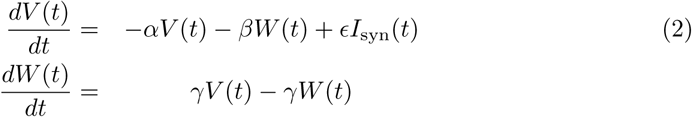

**Table 1.** Parameter descriptions and canonical values used throughout this study, unless otherwise stated. The values of these parameters correspond to an intrinsic subthreshold oscillation period of 31.4 ms.

| Description | Parameter symbol and value |
| --- | --- |
| membrane capacitance | $C=1$ nF |
| leak conductance | $g=0.1$ μS |
| conductance associated with the $w$ variable | $g_w=0.4$ μS |
| time constant of the $w$ variable | $\tau_w=10$ ms |
| threshold voltage | $V_{\text{thr}} = 5$ mV |
| reset voltage | $V_{\text{reset}} = -1$ mV |
| refractory period | $t_{\text{refr}}=3$ ms |

where *α* = *g/C* and *β* = *g*_w_/*C* control the effective leak and the coupling between *V* and *W*, respectively, *ϵ* = 1/*C*, and *γ* = 1/*τ*_w_. If *V > V*_th_ a spike is generated and *V* is reset to *V*_reset_ and kept there for a refractory time *t*_refr_.

The analysis for the conductance-based paradigm discussed below uses the model and parameters described in [25].

### Synapse Model

Incoming synaptic inputs are mediated by an excitatory synapse with short-term depression and facilitation, described according to the Tsodyks-Markram (TM) model [6, 26]:

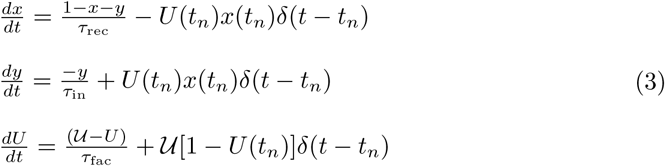

where *x*(*t*) represents the fraction of recovered neurotransmitters, *y*(*t*) the fraction of active neurotransmitters, *U*(*t*) the fraction of released neurotransmitters after the arrival of an action potential, and *t*_*n*_ is the timing of the most recent synaptic event. In this description, *x*(*t*_*n*_) and *U*(*t*_*n*_) are the values of the corresponding variables immediately before the arrival of the most recent synaptic event at *t*_*n*_, and 𝒰 is the fraction of released neurotransmitters at rest. Within this model, the corresponding synaptic current is considered to be proportional to the fraction of active neurotransmitters, i.e., *I*_syn_(*t*) = *𝒜*_syn_*y*(*t*), where *𝒜*_syn_ represents the maximal synaptic strength.

## Results

### Analytical solution of the Generalized Integrate-and-Fire (GIF) neuron with dynamic synapses

In the absence of synaptic inputs, the subthreshold dynamics described by equations (2) is a two-dimensional linear system, whose trajectory from a general initial condition (*V*_0_,*W*_0_) can be expressed as a function of its complex conjugate eigenvalues *λ*_1,2_ = *−μ ± iω*, where 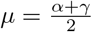 and 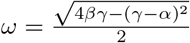. The parameter *μ* determines the membrane time constant, while the parameter *ω* is the oscillation frequency (in rad/ms), related to the intrinsic period of oscillations 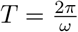 [11].

In the following we consider a synaptic input to the GIF neuron model in the general form of 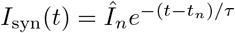 that, as we will see later, has the form of the synaptic current derived from the TM model. With this input, equations (2) constitute a nonhomogeneous differential system, whose solution can be found with the method of undetermined coefficients. That is, we wrote a candidate solution to equations (2) in the form of a sum of an exponential function and the general solution of the corresponding homogeneous system with undetermined coefficients, i.e., we assume for *t > t*_*n*_ the solution:

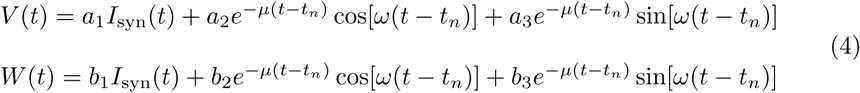

and found the coefficients *a*_*i*_ and *b*_*i*_ that solve (2), resulting in the following values:

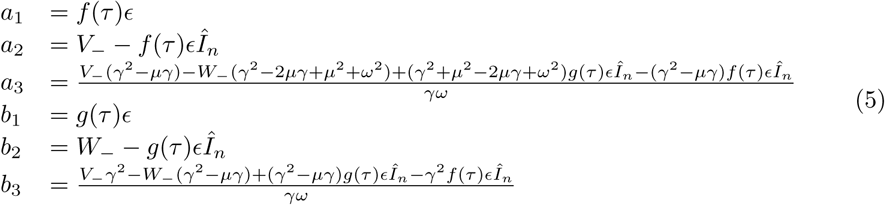

where 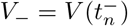, 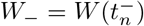 (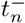 referring to the time just before the input at *t*_*n*_), and

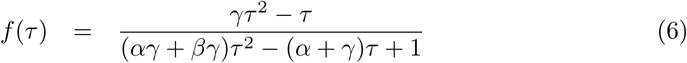

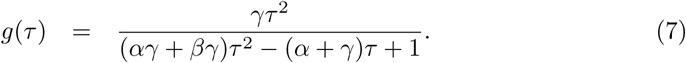

If 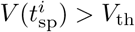 a spike is generated and *V* is reset to *V*_reset_ and kept there for a refractory time *t*_refr_, during which the dynamical variables evolve as

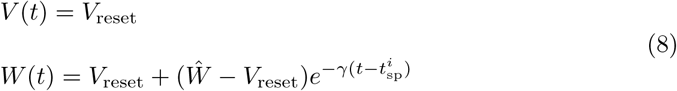

where

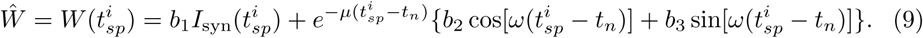

The last term in equation (8) was obtained after integration of the second equation of the system (2) with *V* = *V*_reset_. The above result can be easily generalized for a train of input spikes in the form 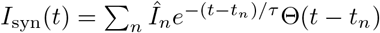.

Note that *f*(*τ*) and *g*(*τ*) scale the synaptic components as a function of *τ* in such a way that for *τ* → 0 (corresponding to infinitely fast synapses) they tend to 0. Hence, in that limit, the terms multiplied by *f*(*τ*) and *g*(*τ*) would vanish.

Between synaptic events, the system (3) that describes the synaptic dynamics is linear and can be easily integrated to obtain

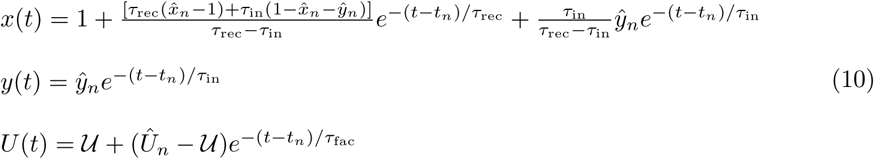

where 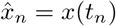, 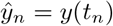 and 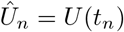. Note that the exponential time dependency of *y*(*t*) guarantees that the synaptic current within this model (which is proportional to *y*(*t*)) meets the functional dependency required above to obtain a general solutions to the GIF neuron model. Moreover, from the system of equations (10), we can derive recursive relations as in [6, 26] to obtain values of 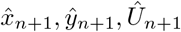 in terms of 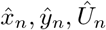. Thus, if we denote ISI=*t*_*n*+1_ *− t*_*n*_ from (10) one obtains for 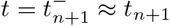

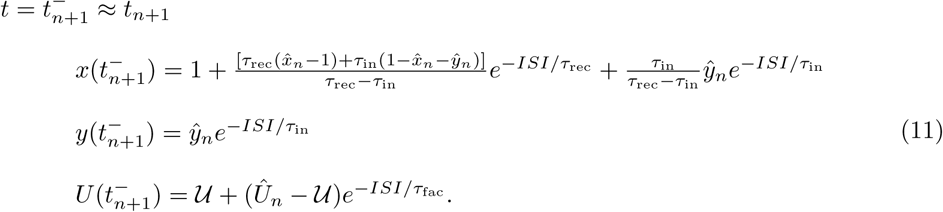

On the other hand, using the original system (3), one obtains

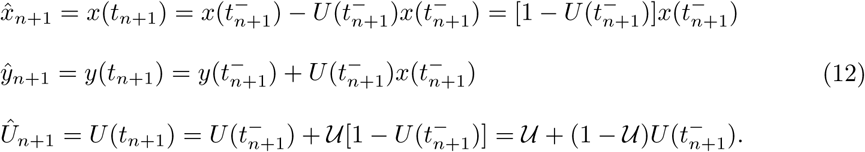

Finally, by substituting (11) into (12), one obtains the recursive relations:

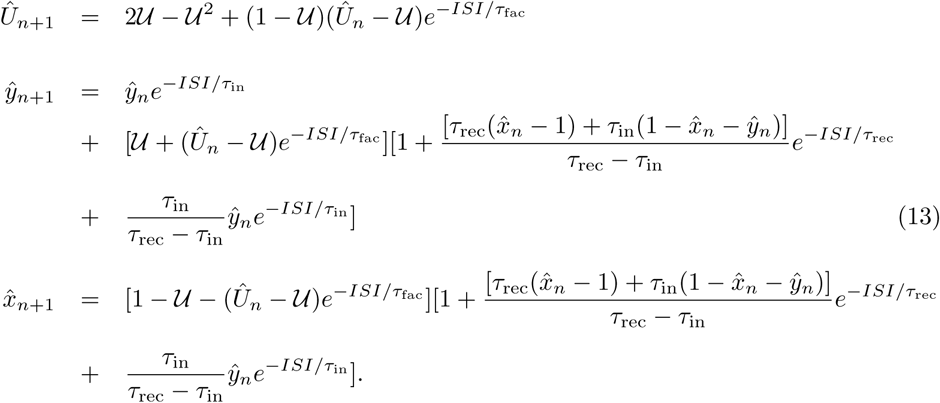

Without loss of generality, initial conditions are set to full availability of neurotransmitters, which corresponds to 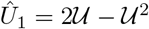, 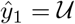 and 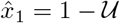. Initial conditions for the neuronal variables are set to the stable point (*V*_0_,*W*_0_)=(0,0). These formulas enable one to compute theoretically the current after each input (*t*_*n*_ *< t < t*_*n*+1_) of a presynaptic spike train as 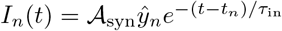 with 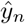 obtained from (13).

Finally, we can use these recursive relations to compute analytically the voltage time series of the GIF neuron when it receives a train of inputs, for both subthreshold and spiking responses. This can be done by setting *τ* = *τ*_in_ and 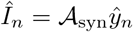 in the general expression for the synaptic current which we used to derive the general solution of the GIF neuron model (see above), i.e., 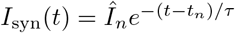 for an input at *t* = *t*_*n*_. Simulations validated that the analytical solution matched exactly the numerical integration of the model. For the analysis discussed below, we will call 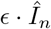 effective postsynaptic potential at the time of the *n* presynaptic spike.

### Nontrivial input-output relationships

The combined action of dynamic synapses and intrinsic neuronal subthreshold oscillations results in a wide variety of input-output transformations from the discrimination of the temporal structure of input trains. To illustrate this, in this paper we focus on nontrivial input-output relationships, such as those characterized by a spike output in response to a decelerating input train and to silence, i.e., a purely subthreshold response, to the corresponding accelerating input train, i.e., to an accelerating input train with the same total duration and constituent ISIs. An integrator neuron without the combined action of its intrinsic oscillations and the dynamic synapse would tend to respond to an accelerating input train and remain silent to a decelerating input. We consider nontrivial responses those cases in which the opposite occurs, and thus the neuron “recognizes” a specific temporal structure in the input pattern. The concept of temporal structure discrimination and nontrivial input-output relationships arising from the combination of intrinsic and synaptic properties is illustrated in Fig. 1. This concept can be generalized in the context of the recognition of any sequence built with a specific temporal structure in the spike train.

**Fig 1.**
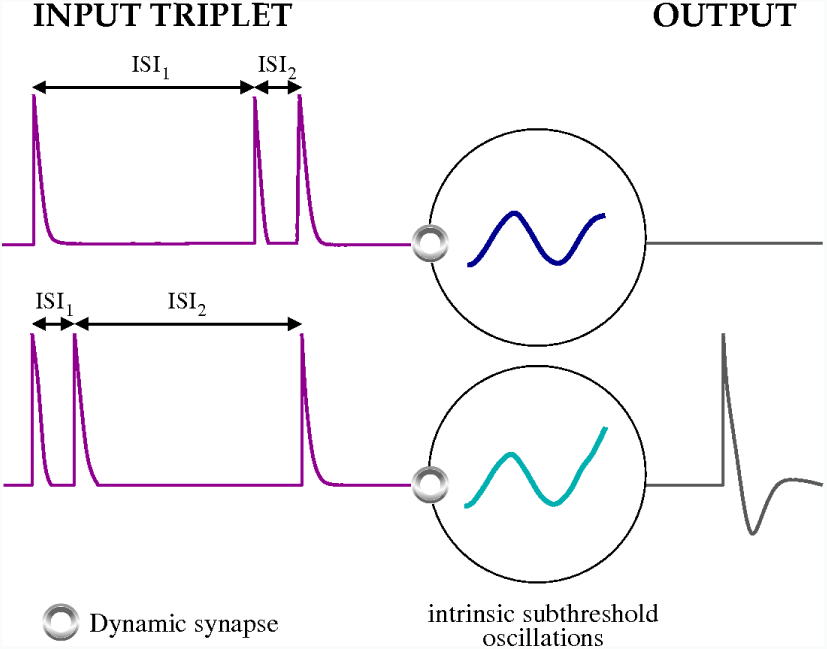
Examples of nontrivial temporal structure discrimination of input signals arising from the combination of intrinsic and synaptic properties. In the top panel, ISI_2_ is shorter than ISI_1_ and yet no response is generated. In the bottom panel, ISI_2_ is longer than ISI_1_ and a response spike is generated after the last spike in the input train. Note that the two input triplets are composed of the same ISIs but in reversed order.

For simplicity, in our analysis we will consider input trains composed of three action potentials (triplets) whose temporal structure can be characterized by the first and second interspike intervals (labeled as ISI_1_ and ISI_2_, respectively). This choice allows a detailed study of the input-output relationships, which can be conveniently visualized in input-output temporal preference maps as described below (Fig. 4).

We focus on a range of synaptic intensities that are insufficient to elicit a postsynaptic spike in response to a single isolated EPSC, but do result in a postsynaptic spike in response to a doublet of incoming EPSCs with appropriate ISI, i.e., either a short ISI or an ISI sufficiently close to the period of subthreshold oscillations or to the period of optimal synaptic facilitation. Figure 2 displays the trajectories of intrinsic and synaptic variables from system (2-3) in nontrivial responses to representative input triplets. These examples illustrate a case where a decelerating input triplet elicits a spike response after the third EPSC, while an accelerating input triplet with the same total duration and constituent ISIs only elicits a subthreshold response. Note that *x* recovers with time constant *τ*_rec_ to model recovery from depression. *U* slowly decays with time constant *τ*_fac_ between neurotransmitter release events to account for facilitation dynamics. The amount of neurotransmitters released in response to a presynaptic event is given by the product *x* ⋅ *𝒰*, which interacts with the state of the neuron to determine the postsynaptic response.

**Fig 2.**
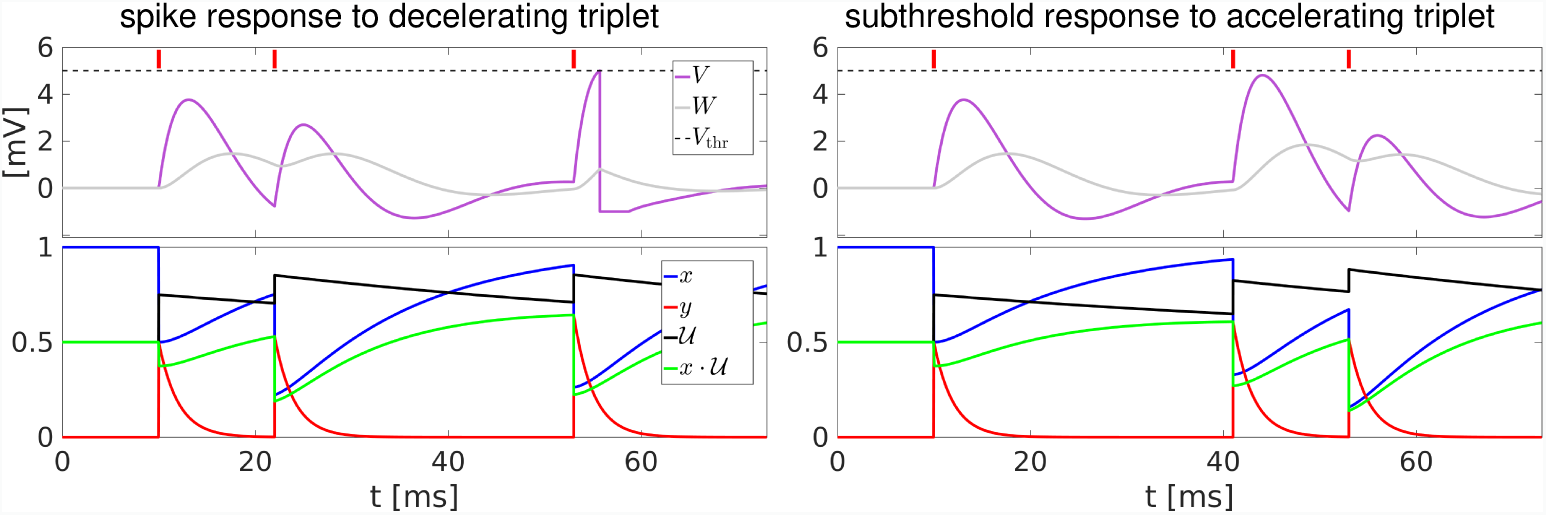
Trajectories of voltage and synaptic variables in response to representative input triplets. Left column, nontrivial response in the form of a spike after receiving a decelerating input triplet (ISI_2_ *>*ISI_1_). Right column, nontrivial silence after receiving an accelerating input triplet (ISI_2_ *<*ISI_1_). Model parameters are specified in Table 1. Synaptic parameters are: *𝒜*_syn_ = 6.3nA, *𝒰* = 0.5, *τ*_rec_ = 14ms, *τ*_fac_ = 60ms, *τ*_in_ = 2ms. Input ISIs are ISI_1_ = 12ms, ISI_2_ = 31ms for the decelerating triplet and ISI_1_ = 31ms, ISI_2_ = 12ms for the accelerating triplet. Red tics indicate EPSC arrival times. Dashed lines indicate the spiking threshold *V*_thr_ = 5*mV*.

In the synaptic model considered here, the synaptic efficacy in response to a given input spike depends on the history of previous stimulation in a complex manner; however, the steady-state synaptic efficacy in response to a train of presynaptic spikes with constant ISI can be easily computed (see [27] and S1 Appendix) as:

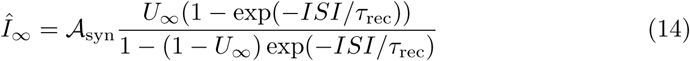

where 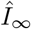 is the steady-state value of 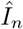 in the limit of *τ*_in_ *→* 0 and

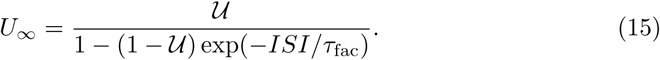

The interaction of synaptic depression and facilitation results in a non-monotonic profile of synaptic efficacies as a function of the input period (Fig. 3), with a peak at the input period that corresponds to maximal synaptic efficacy, which we define as the input period of optimal synaptic facilitation (triangles in Fig. 3).

**Fig 3.**
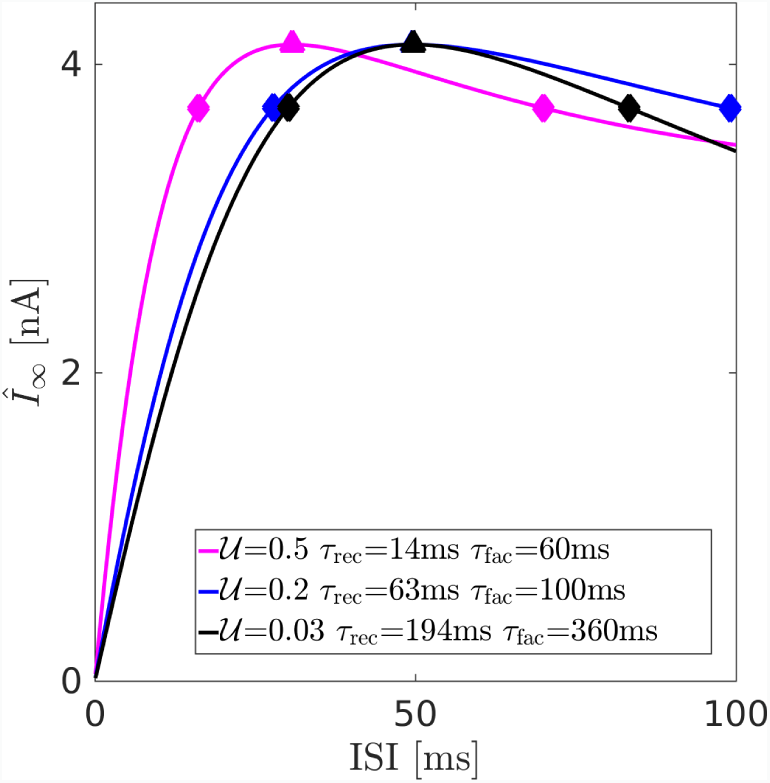
EPSC amplitude as a function of input ISI in the steady-state approximation. The EPSC amplitude as a function of input ISI is shown for three illustrative parameter sets in the steady-state approximation, that is, in response to an infinite train of input spikes with constant ISI. Triangles show peak values and diamonds indicate the points where the EPSC amplitude has fallen to 90% of the peak value for each parameter set. The blue and the black curves have been rescaled to exhibit the same peak value as the magenta curve to facilitate comparison.

In the next sections, we will describe the selective and nontrivial input-output transformations that result from the interplay of intrinsic and synaptic preferences in these three representative cases, and compare them with the corresponding transformations performed by the same GIF neuron but with incoming spike trains delivered through a static synapse.

### Temporal discrimination maps

Using the analytical solution reported above, we can easily calculate the response of the neuron to different input triplets (ISI_1_,ISI_2_) delivered through the dynamic synapse. The use of spike triplets facilitates the representation of the neuron response to multiple input temporal structures in what we call temporal discrimination maps (see Fig. 4). In this map bidimensional representation, the *x*-axis corresponds to ISI_1_ and the *y*-axis to ISI_2_ duration. The color code indicates when and how spikes are produced in response to the incoming input triplet with a specific temporal structure. In these maps, nontrivial input-output responses can be readily identified, as well as the overall dependence of the synaptic parameters in shaping the temporal structure discrimination.

**Fig 4.**
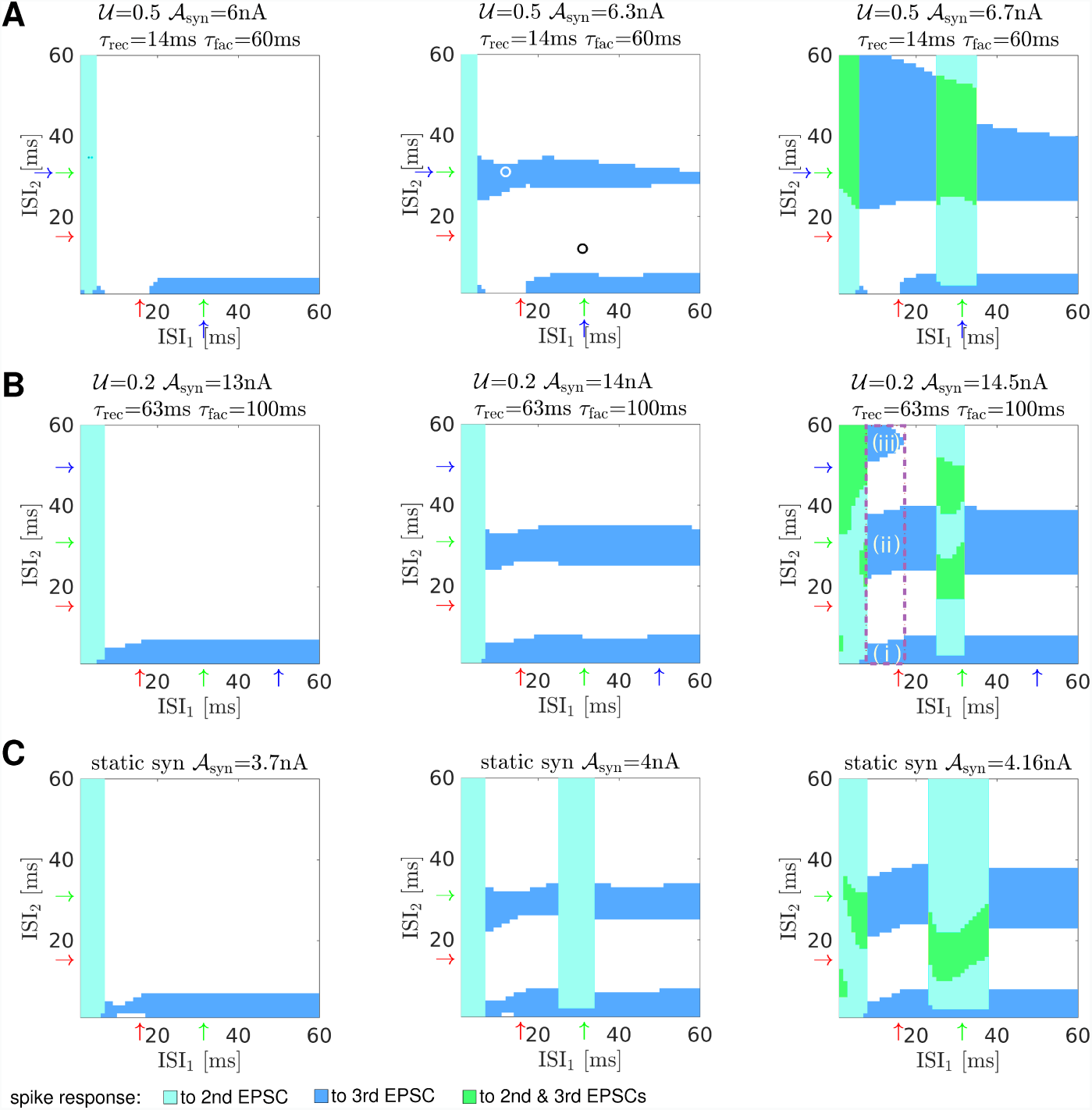
Input-output temporal preference maps. Spike responses to input triplets as a function of the first and second interspike intervals for several representative cases. Arrows indicate the preferred intrinsic ISI, equal to the period of intrinsic oscillations (green); the anti-preferred intrinsic ISI, equal to half the period of the intrinsic oscillations (red); and the preferred synaptic ISI according to the steady-state approximation (blue). A: coincident intrinsic and synaptic preference. B: incommensurate intrinsic and synaptic preference. C: static synapse. The white and black circles in panel A, middle, indicate the input triplets illustrated in Fig. 2, left and right panels respectively. In panel A and B maps *τ*_in_ = 2*ms*. The dashed magenta rectangle and the letter labels in the rightmost map in panel B refer to the explanation of Fig. 5.

Each row in Fig. 4 illustrates three representative cases of distinct intrinsic and synaptic co-action building up the neuronal response as a function of different synaptic strength *𝒜*_syn_, from a low to a moderate-high value (cf. left, middle and right panels). The top row shows temporal discrimination maps for coincident intrinsic and synaptic preferences. The middle row displays the maps for incommensurate intrinsic and synaptic preference. Finally, the last row shows the maps for a static synapse. Nontrivial preferences are only observed for strong enough synapses. Note the regions in these maps that correspond to the neuron’s response only after the third EPSC (dark blue), and how the location and size of these regions is shaped by the parameters of the dynamic synapse.

Middle and right maps of Fig. 4A,B provide an explanation for the nontrivial response to decelerating and acelerating triplets in the illustrative example shown in Figs. 1 and 2. If ISI_*s*_ (with subindex *s* indicating the shortest ISI of the triplet) is sufficiently close to half the period of the subthreshold oscillations (the anti-preferred intrinsic ISI, indicated by a red arrow), decelerating triplets (ISI_*s*_,ISI_*l*_) often elicit a spike response after the third EPSC, with the corresponding accelerating triplet (ISI_*l*_,ISI_*s*_) eliciting only subthreshold responses. This can occur if ISI_*l*_ is close to the period of the subthreshold oscillations (the preferred intrinsic ISI, indicated by a green rightward arrow), close to the preferred synaptic ISI as determined by the steady-state approximation and indicated by a rightward blue arrow, e.g., Fig. 4B, right panel, region indicated as (iii), or in a broader range of input ISIs sufficiently longer than the anti-preferred intrinsic ISI (Fig. 4A, right panel). Nontrivial input-output relations are also present in the maps that correspond to the effect of a static synapse on a neuron with subthreshold oscillations (Fig. 4C, middle and right panels). However, they are more common and shapeable in the maps that correspond to dynamic synapses, as well as more robust to changes in input strength and noise, as we will demonstrate in subsection “Robustness of nontrivial input-output relationships to changes in input strength and noise”. The isolation and modulation of these regions depend on the specific parameters of the synaptic channel.

To favor a mechanistic understanding of how intrinsic and synaptic dynamics interact to shape the neuronal input-output temporal response, it is instructive to assess how the membrane potential and the effective postsynaptic potential covary for a range of input ISIs. To this end, we plot in Fig. 5A the distance to threshold *V*_thr_ *− V* against the effective postsynaptic potential (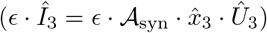) for an illustrative range of (ISI_1_,ISI_2_). This range corresponds to the magenta rectangle in Fig. 4B, right panel. The three (ISI_1_,ISI_2_) disjoint regions shown in yellow, purple and gray correspond to the regions with a spike response to the third EPSC indicated as (i), (ii) and (iii) in Fig. 4B, right panel, respectively.

**Fig 5.**
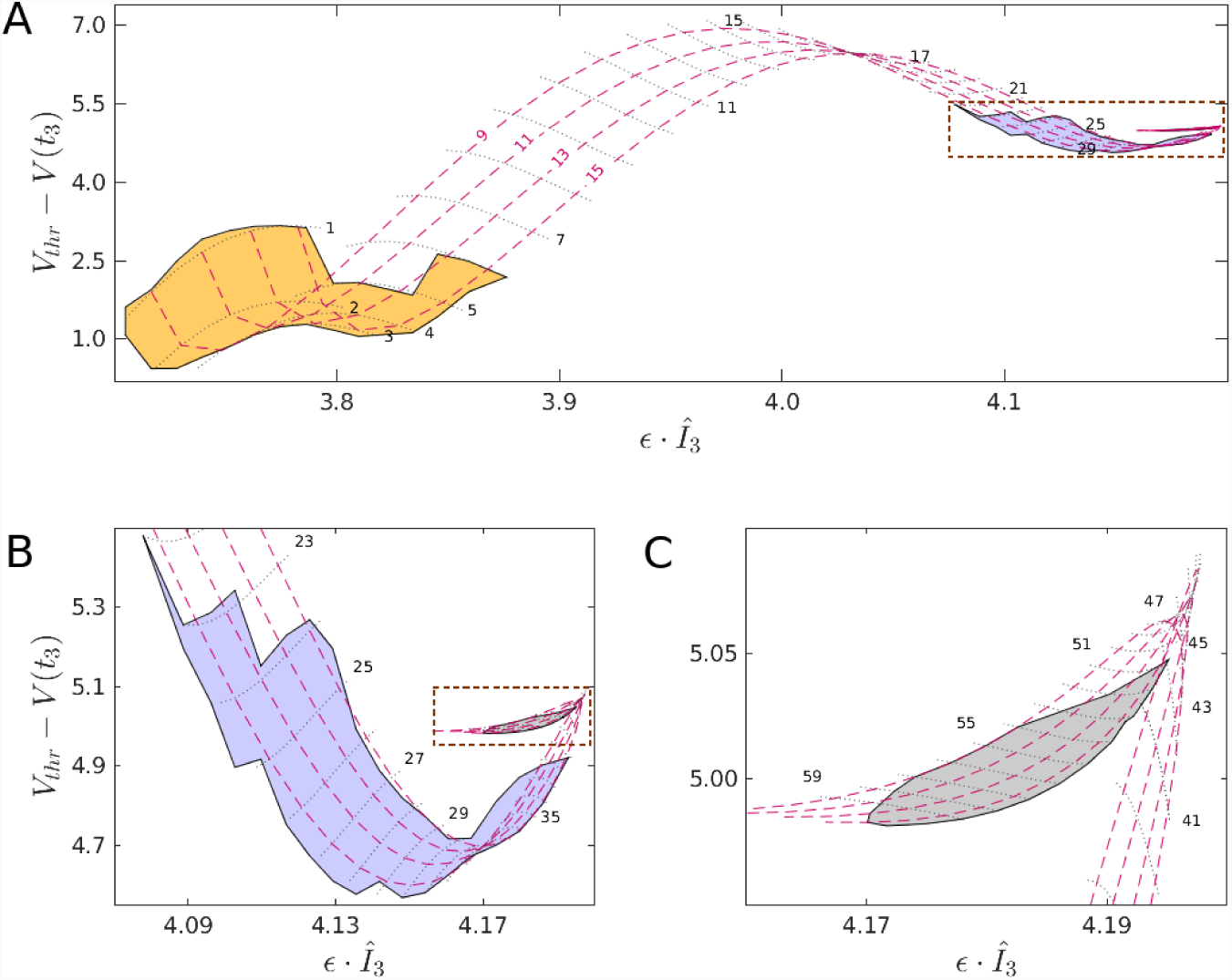
Input-output temporal preferences are shaped by the interaction of intrinsic and synaptic dynamics. The distance to the spike threshold *V*_thr_ *− V* is plotted against the effective postsynaptic potential (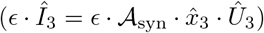) at the time of the third input spike for an illustrative range of (ISI_1_,ISI_2_), which corresponds to the magenta rectangle in Fig. 4B, right panel. In Panels A and B, the dashed rectangle indicates the region that is shown at a larger scale in the following panel. Yellow, purple and gray regions correspond to areas (i), (ii) and (iii) in Fig. 4B, right panel, respectively. Red dashed isolines represent ISI_1_ duration, while black dotted isolines represent ISI_2_ duration.

Depending on the current state of the subthreshold dynamics and on the effective postsynaptic potential resulting from the synaptic dynamics, we observe in Fig. 5 different trade-offs that give rise to the temporal discrimination. The contribution of synaptic dynamics is especially conspicuous for long values of ISI_2_, where intrinsic oscillations have nearly completely waned; conversely, synaptic dynamics exert a mostly modulatory role for shorter values of ISI_2_, where the neuronal response is dominated by the faster intrinsic dynamics. The yellow region in Fig. 5A corresponds to a short distance to the firing threshold. The spike response here is due to the non-instantaneous kinetics of the synaptic pulse resulting from the second EPSC. This type of response is also observed in a similar range of (ISI_1_,ISI_2_) for the GIF model with a static synapse (Fig. 4C). The purple region corresponds to a larger distance to the threshold. Here, the input-output preference is mainly due to the resonant dynamics provided by the restorative current *I*_w_. The synaptic dynamics modulate the extent of this region; in particular, it extends the spike response to longer values of ISI_2_ as synaptic facilitation keeps increasing during the repolarization phase following the depolarization peak. However, it is also present in a similar (ISI_1_,ISI_2_) range for the GIF neuron with static synapse (Fig. 4C). Finally, the gray region is of particular interest, since it clearly emerges as a qualitatively novel phenomenon from the interaction of intrinsic and synaptic dynamics. It is located in an area of strong synaptic facilitation; however, it does not correspond to the peak of synaptic facilitation, but occurs for slightly longer values of ISI_2_, which correspond to a more favorable phase of the intrinsic subthreshold oscillation.

Similar results to those shown in the maps of Fig. 4 were obtained with a GIF neuron with slower intrinsic oscillations (intrinsic period equal 50 ms) and the same *μ/ω* ratio (Fig. 6). However, in this case we did not observe a region of nontrivial preference corresponding to the preferred synaptic ISI at 31.4 ms, probably because of its proximity to the anti-preferred intrinsic ISI (25 ms, Fig. 6A). Conversely, in the case of an integrate-and-fire (IF) neuron with a purely passive subthreshold dynamics, short-term synaptic dynamics modulated input-output relationships but did not result in nontrivial relationships of the kind we focus on here (i.e., one spike response to a decelerating triplet after the third EPSC, silence to the corresponding accelerating triplet of the same total duration and constituent ISIs) (Fig. S1). This highlights the key role played by intrinsic suppression of postsynaptic responses for anti-preferred input ISIs due to intrinsic subthreshold oscillations in the emergence of this kind of nontrivial preference. Note, however, that nontrivial two spike responses can also be observed in the IF model (two spike response to a decelerating triplet, with only one spike in response to the corresponding accelerating triplet), even when inputs are delivered through a static synapse, due to after-spike refractoriness. The location of these nontrivial two spike responses along the ISI_2_ axis is modulated by the properties of the dynamic synapse (Fig S1, middle column).

**Fig 6.**
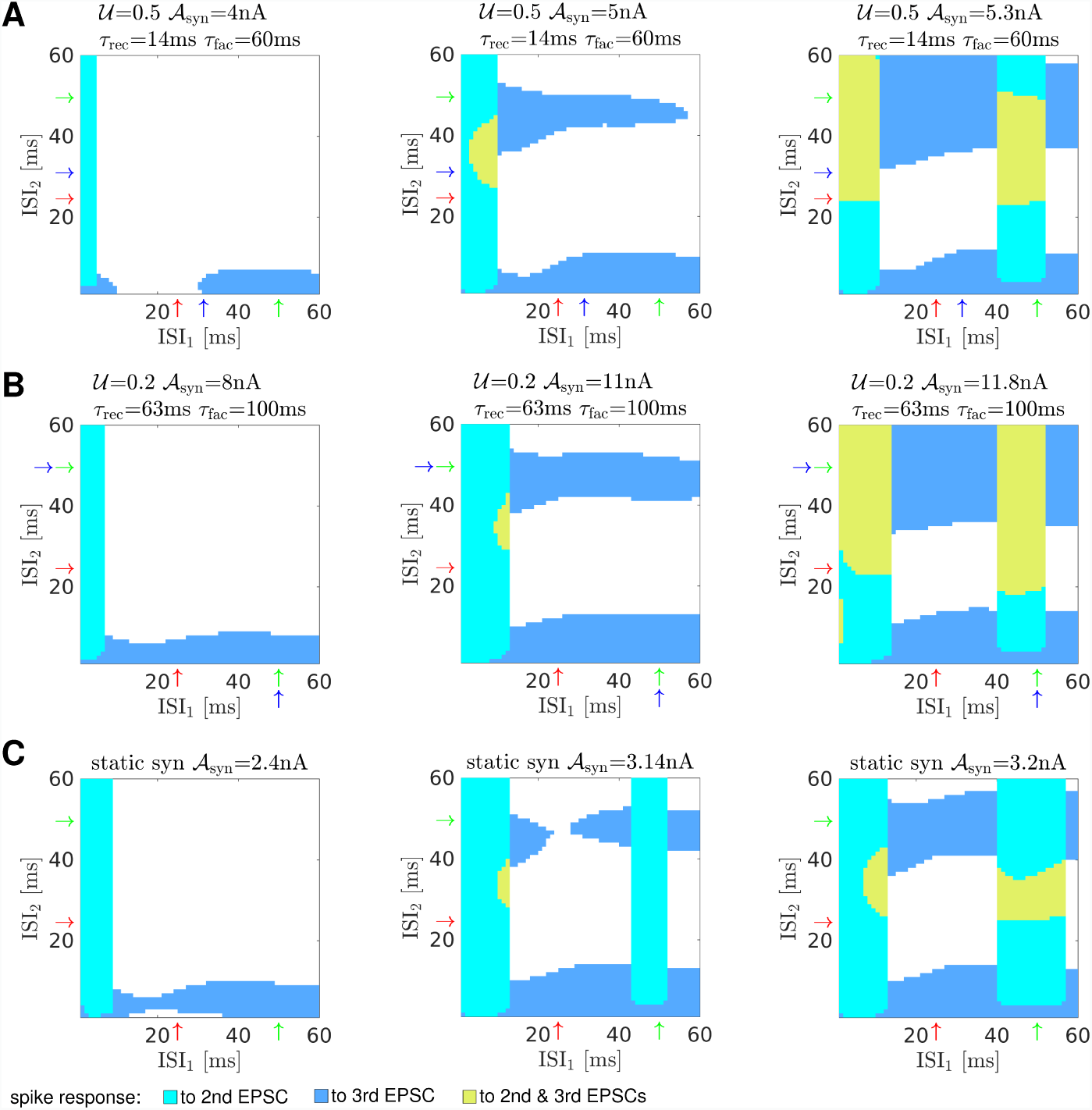
Input-output temporal preference maps for a neuron with slower intrinsic oscillations. As in Fig. 4, for a neuron with slower intrinsic oscillations with period 50 ms. Parameters as in Fig. 4, but *g*=0.0257 *μ*S and *g*_w_=0.1717 *μ*S.

The maps show that the combined effect of the dynamic synapse and the intrinsic oscillations shapes the discrimination of the temporal structure of the input. It is important to emphasize not only the presence of nontrivial preferences in the form of a spike response to decelerating input, but also the silent regions with no response to either accelerating or decelerating input trains. The synergistic action of intrinsic and synaptic properties can lead to the formation of a preference for a very specific temporal structure in the input.

### Robustness of nontrivial input-output relationships to changes in input strength and noise

The interaction of subthreshold oscillations and dynamic synapses not only results in broader regions of the input space (ISI_1_,ISI_2_) where nontrivial input-output preferences are observed, but also in nontrivial preferences which are more robust to parameter changes or noise than what observed with static synapses.

To estimate the degree of robustness of the nontrivial preferences to changes in the maximal synaptic strength *𝒜*_syn_, we performed a series of calculations to determine the “depth” of nontrivial preferences along the direction determined by the maximal synaptic strength *𝒜*_syn_. For each value of (ISI_*s*_,ISI_*l*_), with ISI_*s*_ < ISI_*l*_, which resulted in a nontrivial preference for a given model and *𝒜*_syn_ value, we gradually increased *𝒜*_syn_ up to the point where the input-output transformation ceased to be nontrivial (for example, because both the decelerating and the accelerating triplet would result in a single action potential in response to the third EPSC). We defined *𝒜*_syn,H_ as the highest maximal synaptic strength where a nontrivial preference is observed, such that 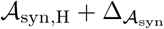 would result in a trivial preference (with 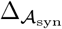 indicating the resolution of the analysis, set to 0.001 nA). Analogously, we gradually decreased *𝒜*_syn_ to the point where the input-output transformation ceased to be nontrivial (for example, because neither the decelerating nor the accelerating triplet would result in any action potential), and defined *𝒜*_syn,L_ as the lowest maximal synaptic strength where a nontrivial preference is observed, also determined with resolution 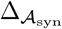.

Then, we defined the percentual depth of nontrivial preference for each value of (ISI_*s*_,ISI_*l*_) that results in a nontrivial preference for a given model and *𝒜*_syn_ as (*𝒜*_syn,H_ *− 𝒜*_syn,L_)/*𝒜*_syn_. The percentual depth of nontrivial preference for the three models of dynamic synapses introduced in Fig. 3 and a model of static synapse are shown in Fig. 7.

**Fig 7.**
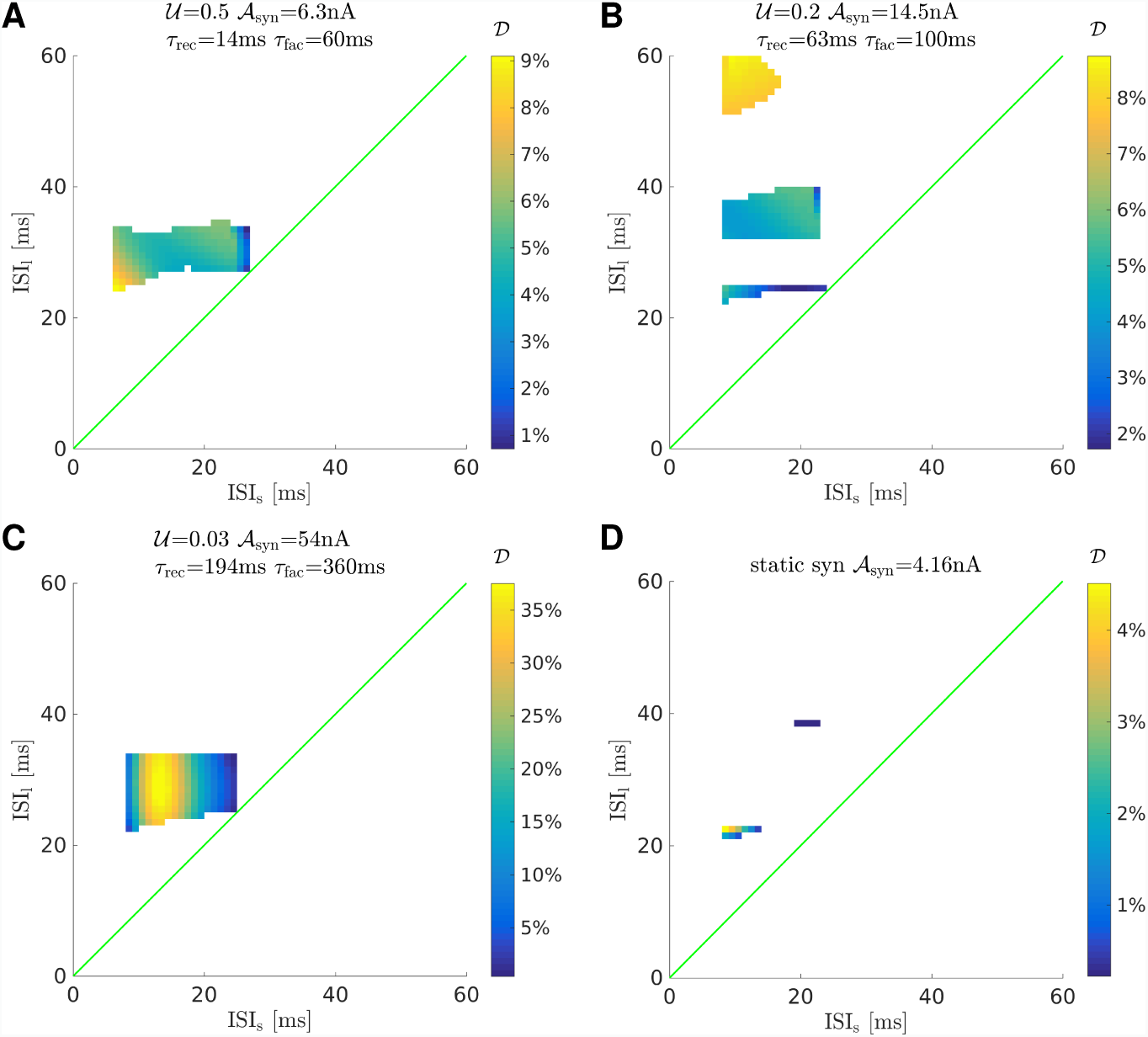
Depth of nontrivial preference relationships for several representative parameter sets. A: coincident intrinsic and synaptic preference. B: incommensurate intrinsic and synaptic preference. C: incommensurate intrinsic and synaptic preference, slow time constants. D: static synapse.

For coincident intrinsic and synaptic preferences (Fig. 7A), nontrivial preferences are observed for a broad region of the input space (ISI_*s*_,ISI_*l*_), with ISI_*l*_ close to the period of the intrinsic subthreshold oscillations and the optimal synaptic facilitation (which are equal by construction for this parameter set). For incommensurate intrinsic and synaptic preferences (Fig. 7B), an additional region of nontrivial preferences is observed for longer ISI_*l*_, close to the period of optimal synaptic facilitation (equal to 50 ms according to the steady-state approximation for this parameter set). The depth of nontrivial preferences can reach values of 8% or 9% for these two models, and can increase above 35% if longer time constants for synaptic depression and facilitation are considered (Fig. 7C). Conversely, for the GIF neuron with a static synapse, only a few points in the input space (ISI_*s*_,ISI_*l*_) correspond to nontrivial preferences (Fig. 7D), and these are very shallow in the *𝒜*_syn_ direction, reaching a maximum depth of only 4.5%.

We also assessed the robustness to noise of the combined action of dynamic synapses and intrinsic subthreshold oscillations in shaping the recognition of specific temporal structure in the input signals. We included Gaussian noise as an additive term in equation 2 as *η* = *𝒜*_noise_ *⋅ 𝒩* (0, 1), being *𝒜*_noise_ a scale factor for the amplitude noise, and simulated 100 independent realizations for each input triplet (ISI_1_,ISI_2_) with a 0.005*ms* time step. Movies S2 to S10 show probabilistic temporal preference maps in the presence of different noise levels. For each input triplet and noise level, the color code from the deterministic temporal preference maps was used if the corresponding output was observed in at least 80% of the simulations, otherwise it was indicated in gray. Note that the nontrivial input-output relationships are preserved for low (corresponding to oscillation amplitudes in response to the noise alone up to 0.15 mV) and moderate levels of noise (up to 0.27 mV), and are only destroyed for a high noise level (up to 0.70 mV).

### Generalization of the synergistic phenomena in a conductance-based model

To test the generalization of the results discussed for the GIF model in a more biophysical description, we have used a Hodgkin-Huxley type formalism proposed to model subthreshold oscillations of inferior olive (IO) neurons [25] and a current modeling synaptic input through a dynamic synapse model. The single-compartment cell model consists of five voltage-dependent ionic currents – a sodium current (*I*_*Na*_), a persistent sodium current, (*I*_*Nap*_), a potassium delayed rectifier current (*I*_*Kd*_), a slow inactivating potassium current (*I*_*Ks*_) and a hyperpolarizing potassium current (*I*_*h*_) – and a leakage current (*I*_*l*_). The neuron model generates characteristic subthreshold oscillations and spiking activity in the amplitude and frequency ranges observed in the living IO cells, which implies slower dynamics than the GIF model discussed above. A detailed description of the model, its parameters and behavior can be found in [25].

We have previously used this neuron model to study the interplay between subthreshold oscillations and a dynamic synapse with a depressing mechanism in response to bursting input [15]. In this section, the synaptic input arriving at the conductance-based neuron is described by combining the Destexhe *et al*. model for synaptic conductances [28] and the Tsodyks-Markram’s description of dynamic synapse currents with both depression and facilitation [6, 26]. Thus, we preserve all relevant short-term depression and facilitation features observed in the Tsodyks-Markram model while providing the dynamic synaptic conductance description required by a conductance-based model. For a detailed discussion on this topic, see [15].

The conductance-based model has richer dynamics shaping the subthreshold oscillations than the GIF model. This endows the IO neuron model with sustained intrinsic subthreshold oscillations in the absence of synaptic input (*σ* = 1.5 and *I*_*inj*_ = 0.345*μA/cm*^2^), unlike the GIF model. Furthermore, the additional time scales of the conductance model participate in shaping the synergistic interaction with the dynamic synapse in the processing of the temporal structure of the input. Figure 8 illustrates that the conductance-based model is able to produce equivalent nontrivial responses to input triplets as the ones produced by the GIF model (cf. Fig 2). In these simulations, we took special care to reproduce in both cases the phase of the subhreshold oscillation when the first spike arrived. Left upper panel in Fig 8 illustrates that a decelarating input triplet evokes a spike response after the last action potential arriving through the dynamic synapse. It is important to emphasize that the third spike does not generate a spike response by itself. Right upper panel shows the absence of a spike response when the same triplet is reversed. Lower panels show the evolution of the variables shaping the synaptic dynamics. The interpretation of the nontrivial input-output relationships is the same as the one provided for the GIF model.

**Fig 8.**
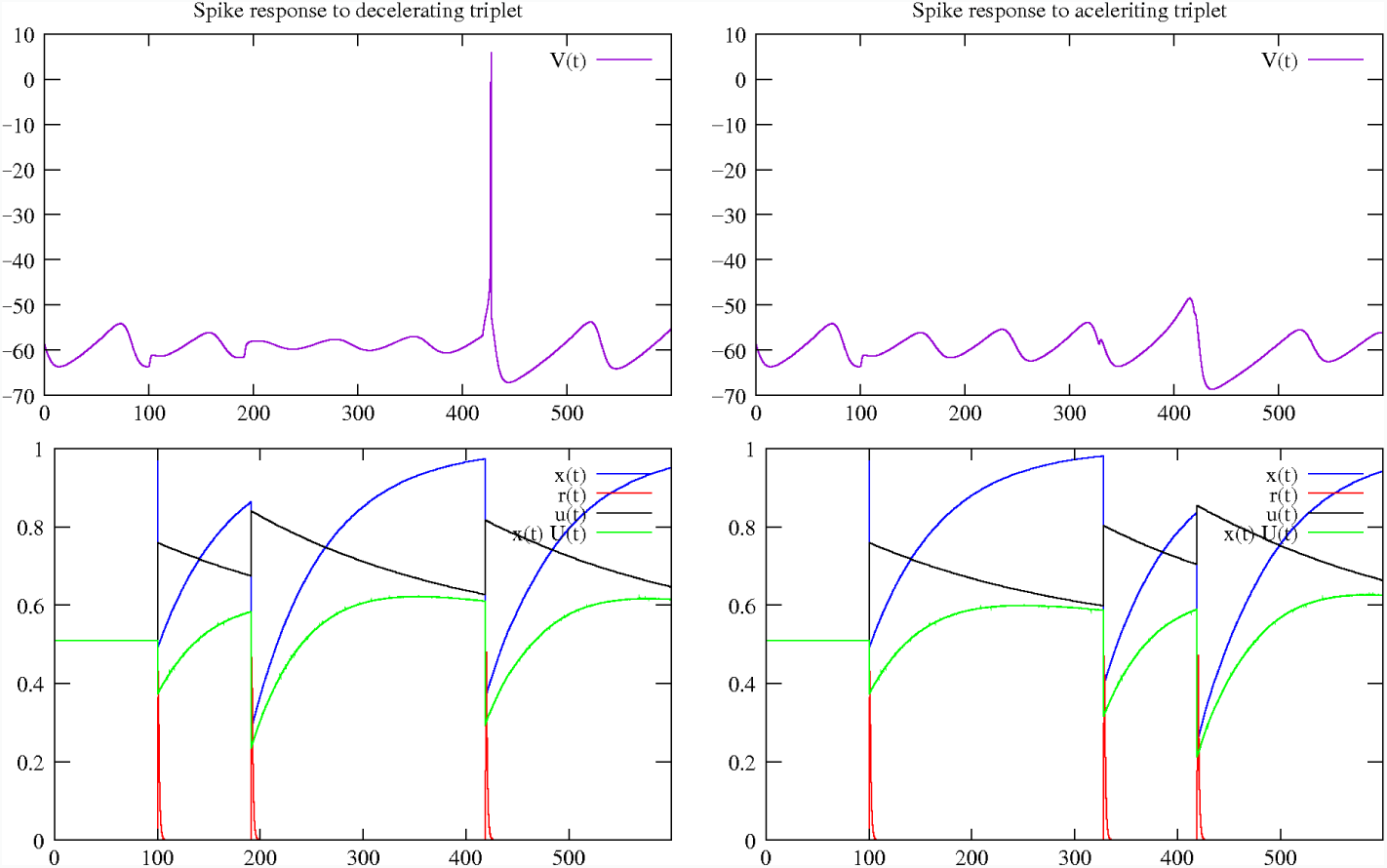
Nontrivial response of a conductance-based model to decelerating and accelerating input triplets. This figure is equivalent to Fig. 2 for the GIF neuron model. Time constants for the dynamic synapse model were adapted to the slower dynamics of the conductance-based model: *𝒰* = 0.51, *τ*_rec_ = 66.5ms, *τ*_fac_ = 211.0ms. Consequently, input ISIs were also longer: ISI_1_ = 91ms and ISI_2_ = 228ms for the decelerating triplet, and ISI_1_ = 228ms and ISI_2_ = 91ms for the accelerating triplet. Results are equivalent as those reported in Fig. 2 for the GIF model.

## Discussion

Intrinsic and synaptic dynamics interact to shape input-output relationships in individual neurons. In this paper, we analyzed the interaction of intrinsic subthreshold oscillations and dynamic synapses with depression and facilitation. We derived an analytical solution for the GIF model in response to inputs delivered through dynamic synapses. Using this solution, we investigated how the combination of intrinsic oscillations and of the input modulation by a dynamic synapse gives rise to preferred and anti-preferred input-output relationships. These temporal preferences do not require fine-tuning of synaptic strength, they are robust to noise, and they are shapeable by the properties of the dynamic synapse in a channel-specific manner. For simplicity, we focused our analysis in the context of input triplets, but our methodology can be readily generalized to more complex input spike trains.

From the reported results, we conclude that the interaction of intrinsic and synaptic properties can enable the implementation of robust channel-specific input discrimination mechanisms for the emergence of selective neuronal responses. These mechanisms are likely to be relevant for decoding temporally precise spiking patterns *in vivo* given the experimental evidence for target-specific synaptic dynamics [29] and tuning of synaptic integration [14]. Recent experimental results have also associated synaptic diversity with the processing of temporally precise multisensory information at the level of individual cells [20].

Single-channel/single-neuron temporal input discrimination can be a computationally economic and metabolically efficient mechanism of contextual information processing with high sensitivity to the precise temporal structure of spiking activity in heterogeneous networks. The intrinsic dynamics of a single neuron can result in very distinct responses to spike input trains depending on the specific properties of the synaptic channel through which they are delivered.

Our model results provide testable hypotheses regarding the presence of nontrivial input-output preferences that can be addressed in experiments with dynamic clamp in living neurons. In this type of experiments, several alternatives are possible with the implementation of artificial dynamic synapses and intrinsic conductances which allow the exploration of the influence of the corresponding synaptic dynamics and intrinsic or induced subthreshold oscillations on neuronal responses. Alternatively, a pattern-clamp approach [30, 31] with paired recordings can be used to reveal input-output preferences implemented by existing biological dynamic synapses through evoking patterned spike trains on the presynaptic neuron while recording from a postsynaptic neuron with subthreshold oscillations.

The temporal discrimination discussed in this paper arises from the combination of synaptic and intrinsic mechanisms. Beyond the context of precise temporal discrimination, this combination can also result in novel phenomena for information processing based on different types of resonance mechanisms such as phase [32] and stochastic resonance [33, 34], including the recently reported inverse stochastic resonance [35–37]. For instance, the discussed modulation mechanisms can give rise to new resonant relationships, which can be relevant in the context of cell recruitment during brain rhythms in health and disease, and also for temporally precise sensory recognition (e.g., visual or auditory) in noisy environments.

Our theoretical results were obtained for a simplified model that enabled analytical treatment, but we also observed the same temporal discrimination mechanism in a conductance-based model. We expect these phenomena to occur also in more biophysically realistic multicompartmental models displaying subthreshold oscillations, particularly in those that are used to relate subthreshold activity to function, e.g., [38], and, most importantly, in living animals. More complex theoretical descriptions for intrinsic subthreshold dynamics, such as those discussed in [32, 39, 40], are likely to result in richer input-output relationships when combined with dynamic synapses. In this paper we focused on single neuron dynamics, but related phenomena can also be analyzed at the network level in addition to the activity shaped by dynamic synapses [12, 41, 42]. The discussed results also point in the direction of non-exclusive neurocentric views of learning [43] based on the interaction between intrinsic and synaptic dynamics.

Our work has specific relevance in the context of recent studies on the effect of precise spiking coding mechanisms on learning processes in olivo-cerebellar circuits [44, 45], where precise timing requirements are related to both function and learning capabilities. In particular, inferior olive neurons, whose subthreshold oscillations are related to motor coordination [46], receive synaptic inputs that modulate their oscillatory activity [47]. Specifically, IO neurons are the target of synapses from the deep cerebellar nuclei which are heterogeneous and often exhibit short-term synaptic dynamics [48].

While we focused on the interaction between intrinsic subthreshold oscillations and short-term synaptic dynamics, previous modelling work suggested that subthreshold oscillations can also have profound effects on the dynamics of long-term plastic changes such as those induced by spike-timing dependent plasticity (STDP) rules [49]. This phenomenon, together with the recent observation of multiple and channel-specific forms of synaptic plasticity in hippocampal and neocortical circuits [50, 51], suggests that the interaction of intrinsic and synaptic short-term dynamics and long-term plasticity with post- and pre-synaptic specificity will be an important topic for future experimental research. This picture is complicated by the presence of heterosynaptic plasticity [52–54], further highlighting the important role of computational models for summarizing disparate experimental evidence and providing formal frameworks towards mechanistic explanations of these phenomena and of the computational role they play in living animals.

An influential theoretical framework proposes that neuronal microcircuits continuously process incoming information in a context-dependent manner across multiple time-scales for flexible and efficient resolution of behavioral demands [55–61], a phenomenon thought to arise from the interaction of intrinsic and synaptic dynamical processes [62–66]. By exposing the temporally precise information processing capabilities afforded by the combination of two key dynamical components of neuronal networks, intrinsic subthreshold oscillations and short-term synaptic dynamics, this work constitutes a step towards a mechanistic understanding of transient and context-dependent information processing in neuronal microcircuits.

## Acknowledgements

This work was supported AEI FEDER grants FIS2017-84256-P (JJT) and PGC2018-095895-B-I00, DPI2015-65833-P (RL & PV).

## S1 Appendix

To derive equations (14) and (15) we have to focus on the system of equations (11), (12) and (13) of the paper. Lets consider *τ*_in_ ≪1 in the second and third equations of the system (13). We obtain then:

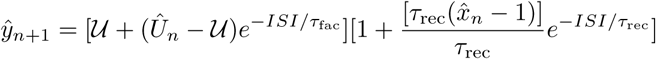

and

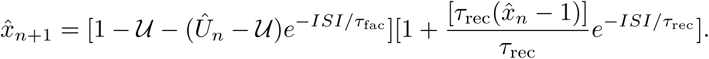

Using here that 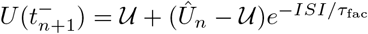 (see last equation of the system (11)) one obtains

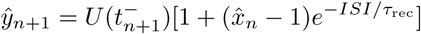

and

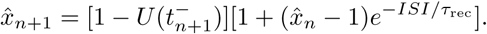

Lets consider now steady state conditions (*n → ∞*) in the last equations and define 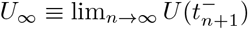. Then, one obtains:

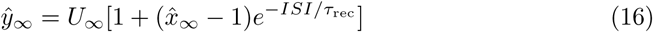

and

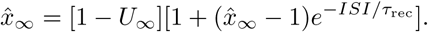

Solving the last equation one obtain:

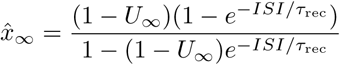

and therefore

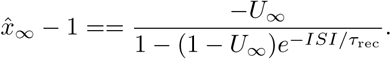

Substituting this last expression in (16) one finally obtains

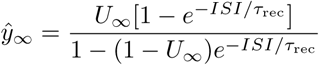

that is, equation (14) in the paper, since 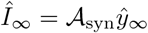.

On the other hand, from the last equation of (11) and the last equation of (12) using the definition of *U*_*∞*_ one obtains

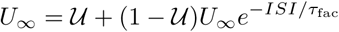

which finally solving for *U*_*∞*_ gives equation (15) in the paper, that is:

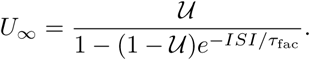

---

**S1 Figure.**
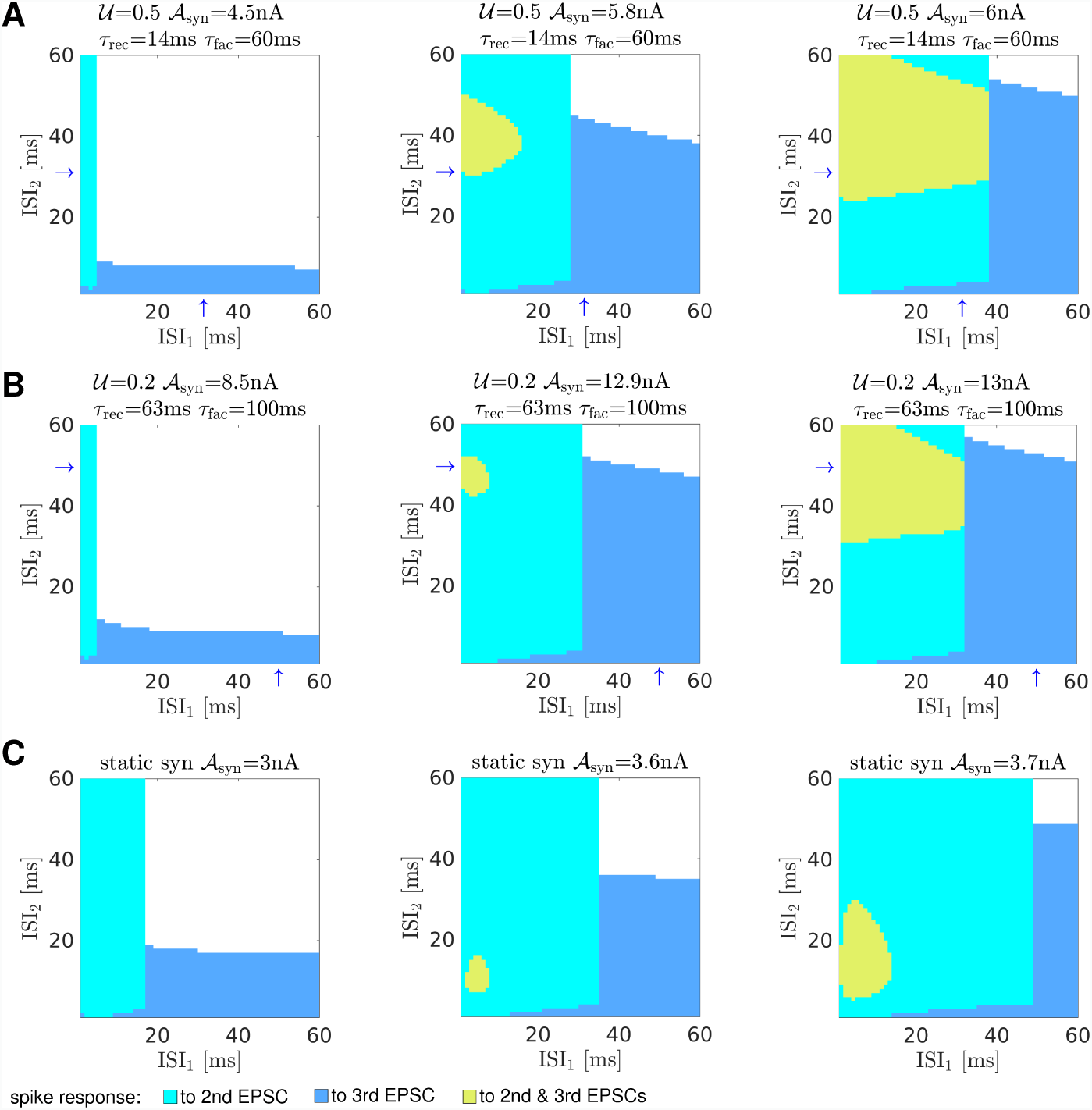
Input-output temporal preference maps for an IF neuron with purely passive intrinsic dynamics. Maps as in Fig. 4, for an IF neuron. Parameters as in Fig. 4, but *g*_w_=0. Blue arrows on panel A and B indicate the preferred synaptic ISI according to the steady-state approximation.

## S1 Movie

**Robustness to noise of the input-output temporal preference map in the left panel of Fig. 4A**. Each frame in the movie corresponds to the discussed map for increasing values of *𝒜*_noise_, from 0 (no noise) to 12.5. Each frame in this movie was generated averaging 100 independent simulations with a statistical threshold of 80%. When an output was observed in at least 80% of the simulations for an input triplet, the corresponding point was plotted with the same color code as in the maps shown in Fig. 4: cyan for an output spike in response to the second EPSC in the input triplet; blue for a spike in response to the third EPSC; and green when the neuron generated a spike in response to both, second and third, EPSCs. When the number of observations for a triplet was below the statistical threshold, the corresponding point in the map was plotted in gray. White indicates no spike response.

## S2 Movie

**Robustness to noise of the input-output temporal preference map in the middle panel of Fig. 4A**. Color codes as explained in S1 Movie.

## S3 Movie

**Robustness to noise of the input-output temporal preference map in the right panel of Fig. 4A**. Color codes as explained in S1 Movie.

## S4 Movie

**Robustness to noise of the input-output temporal preference map in the left panel of Fig. 4B**. Color codes as explained in S1 Movie.

## S5 Movie

**Robustness to noise of the input-output temporal preference map in the middle panel of Fig. 4B**. Color codes as explained in S1 Movie.

## S6 Movie

**Robustness to noise of the input-output temporal preference map in the right panel of Fig. 4B**. Color codes as explained in S1 Movie.

## S7 Movie

**Robustness to noise of the input-output temporal preference map in the left panel of Fig. 4C**. Color codes as explained in S1 Movie.

## S8 Movie

**Robustness to noise of the input-output temporal preference map in the middle panel of Fig. 4C**. Color codes as explained in S1 Movie.

## S9 Movie

**Robustness to noise of the input-output temporal preference map in the right panel of Fig. 4C**. Color codes as explained in S1 Movie.

