## Supplementary figures and images for "Temporal discrimination from the interaction between dynamic synapses and intrinsic subthreshold oscillations"

### S1 Movie

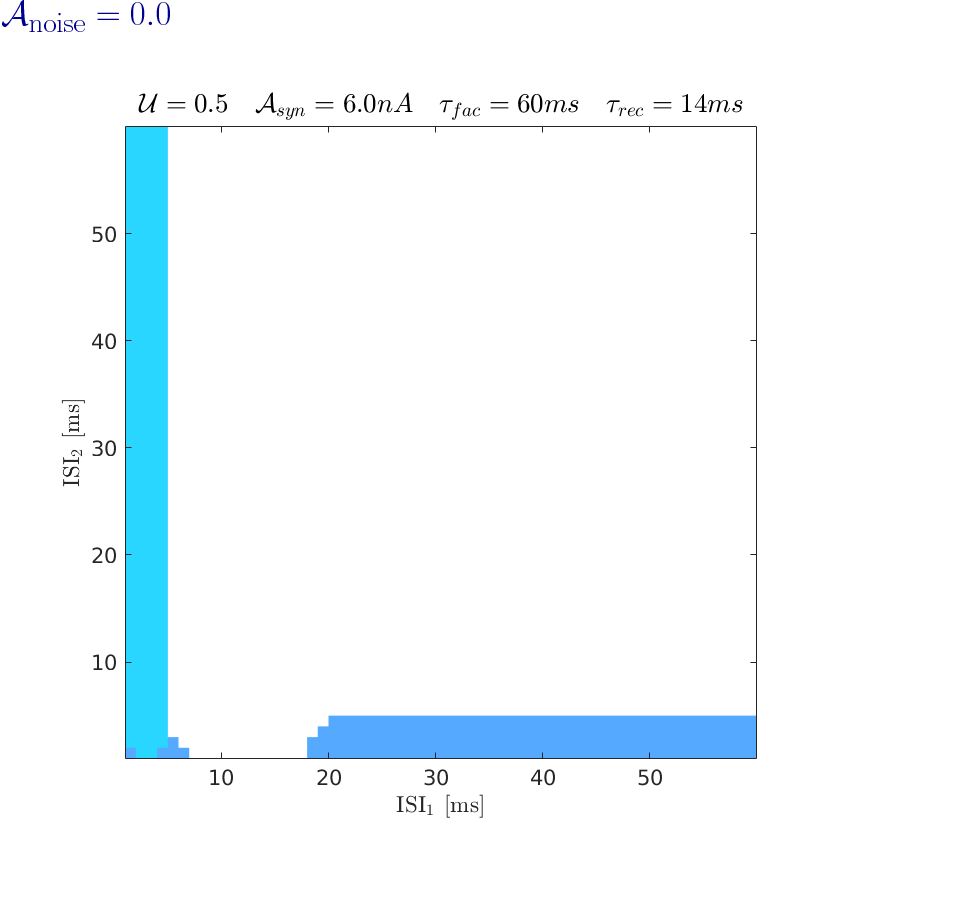

### S2 Movie

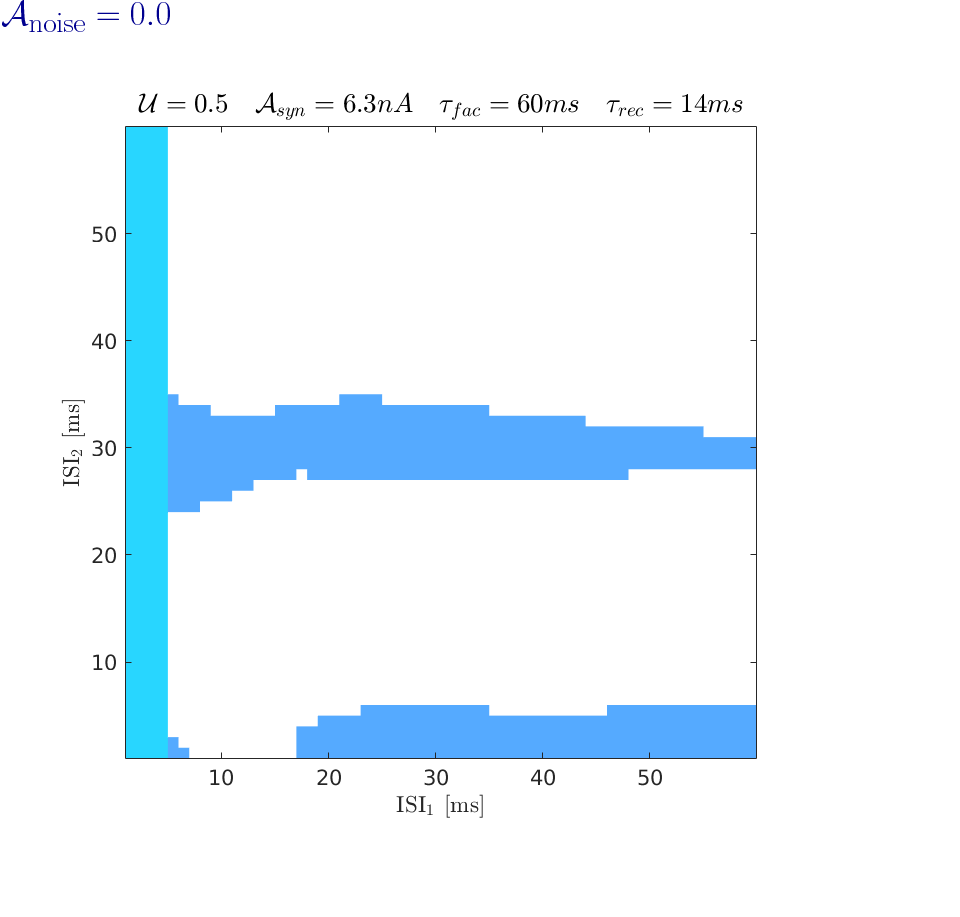

### S3 Movie

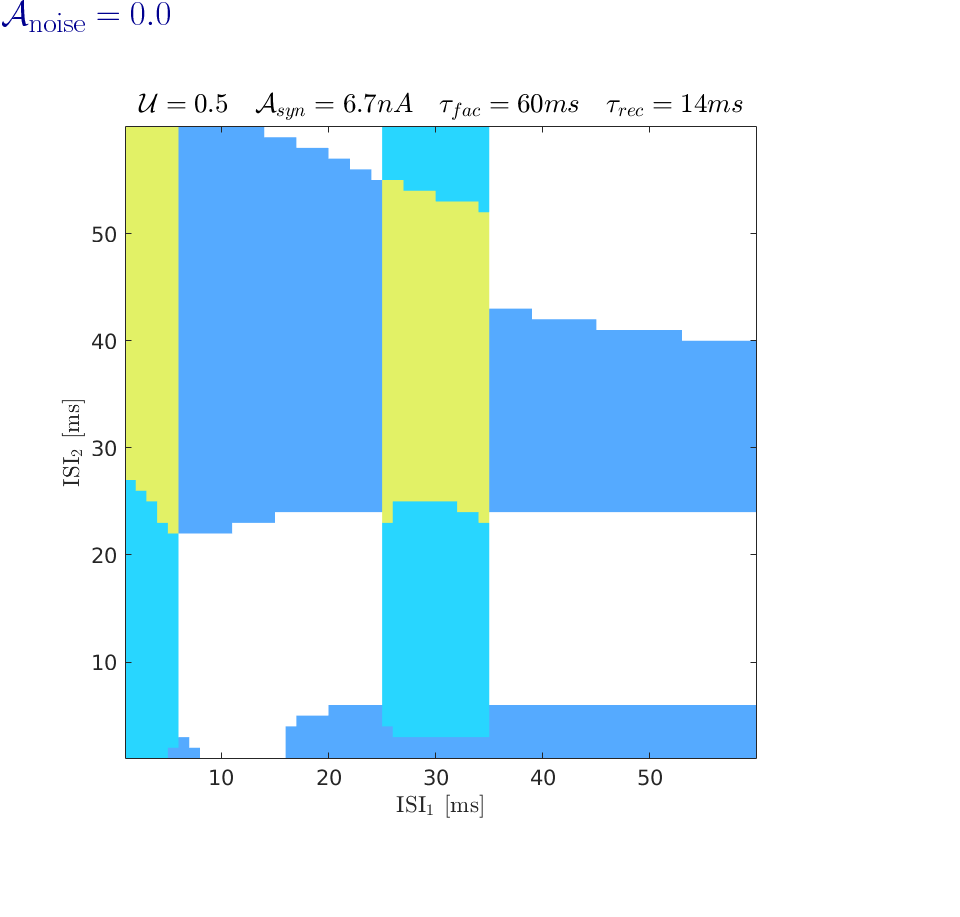

### S4 Movie

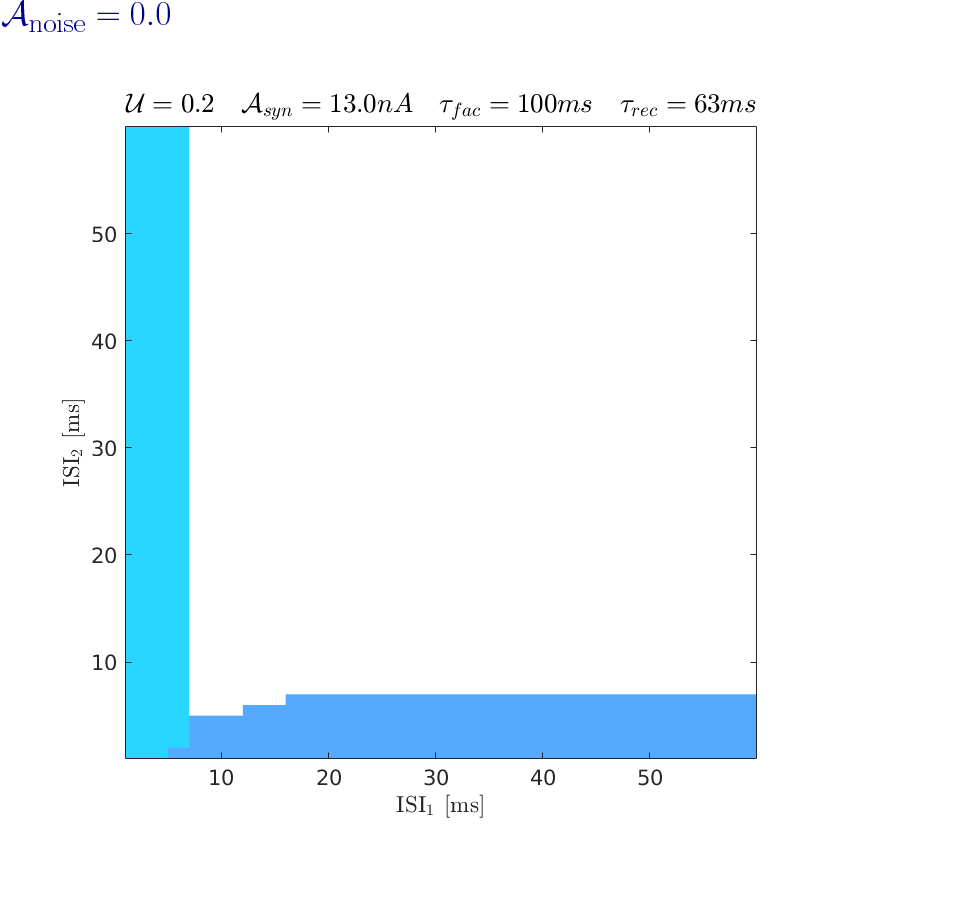

### S5 Movie

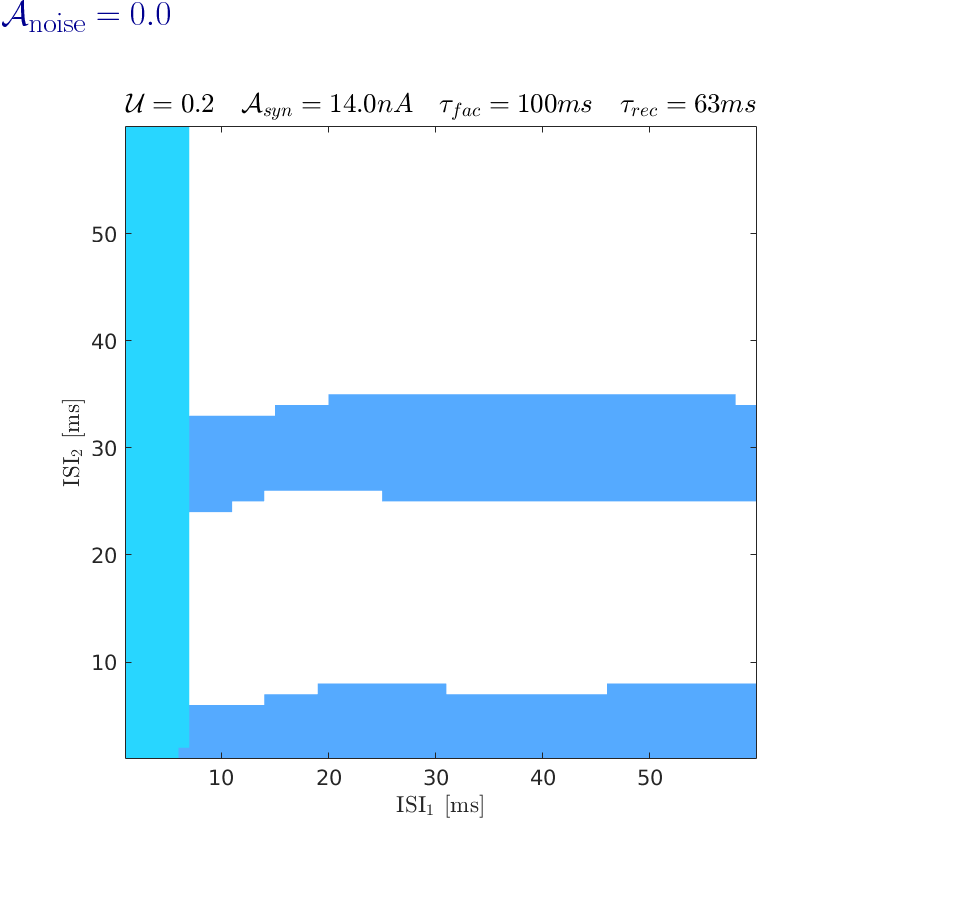

### S6 Movie

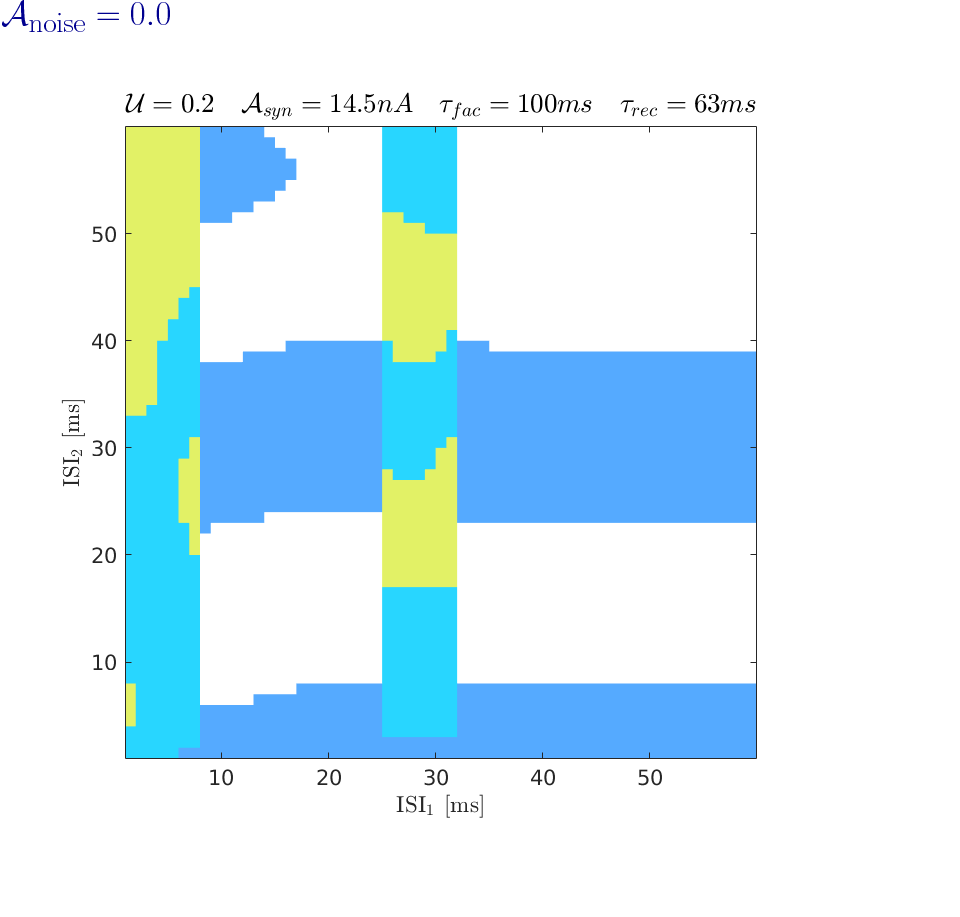

### S7 Movie

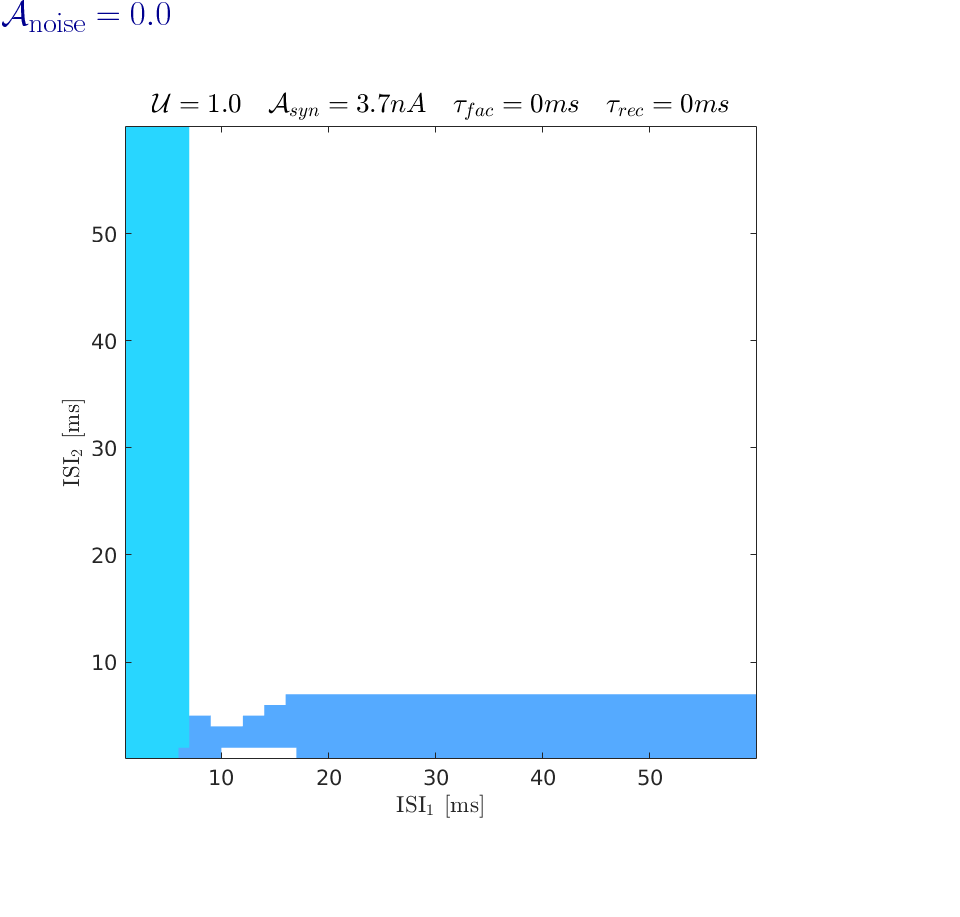

### S8 Movie

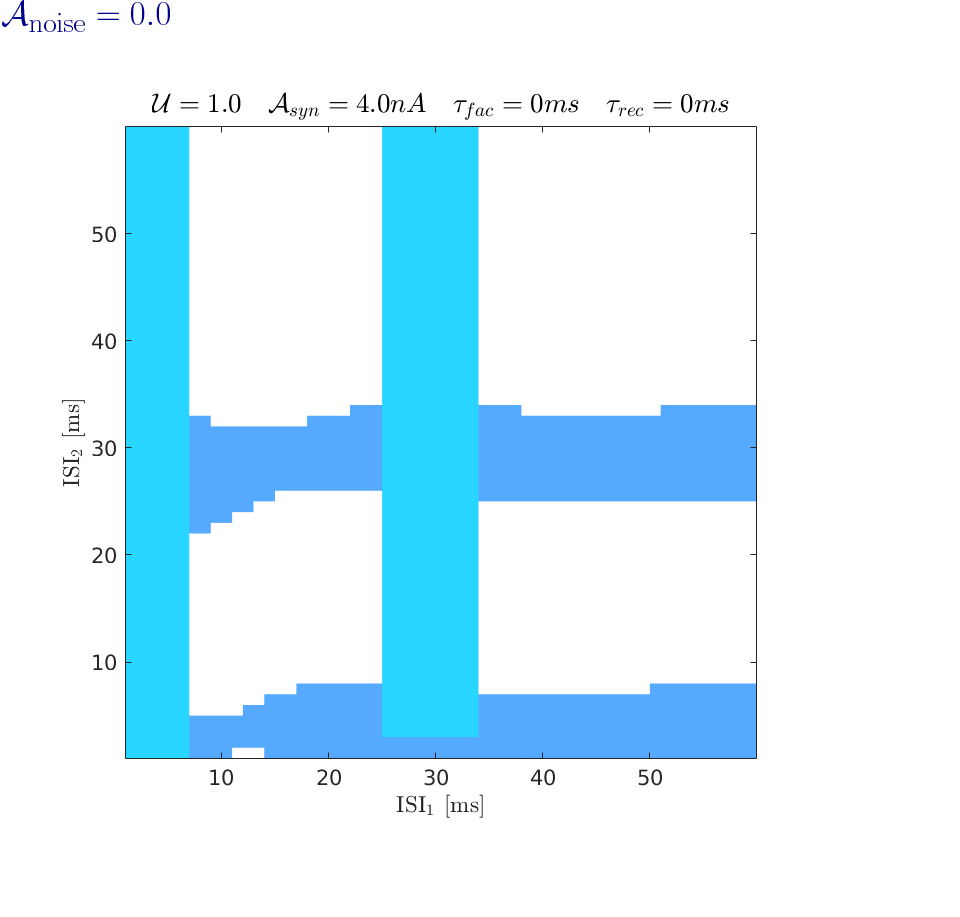

### S9 Movie

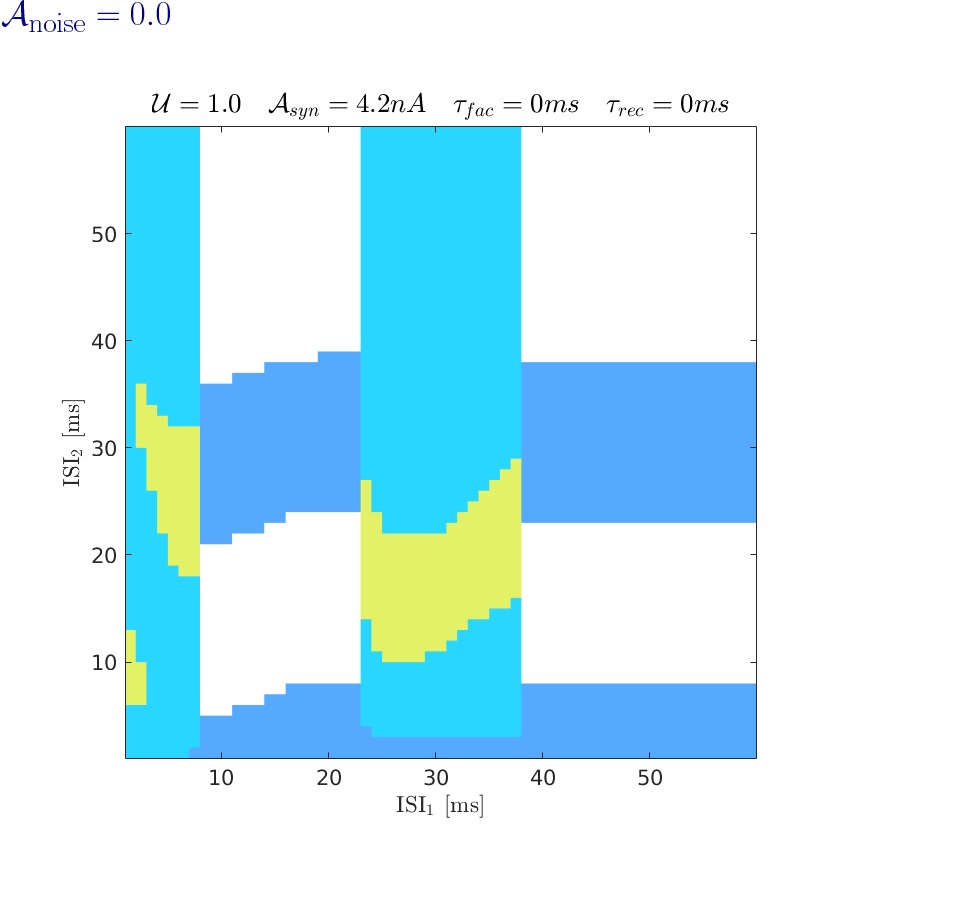
